# *Airpart*: Interpretable statistical models for analyzing allelic imbalance in single-cell datasets

**DOI:** 10.1101/2021.10.15.464546

**Authors:** Wancen Mu, Hirak Sarkar, Avi Srivastava, Kwangbom Choi, Rob Patro, Michael I. Love

## Abstract

**Motivation:** Allelic expression analysis aids in detection of *cis*-regulatory mechanisms of genetic variation which produce allelic imbalance (AI) in heterozygotes. Measuring AI in bulk data lacking time or spatial resolution has the limitation that cell-type-specific (CTS), spatial-, or time-dependent AI signals may be dampened or not detected.

**Results:** We introduce a statistical method *airpart* for identifying differential CTS AI from single-cell RNA-sequencing (scRNA-seq) data, or other spatially- or time-resolved datasets. *airpart* outputs discrete partitions of data, pointing to groups of genes and cells under common mechanisms of cis-genetic regulation. In order to account for low counts in single-cell data, our method uses a Generalized Fused Lasso with Binomial likelihood for partitioning groups of cells by AI signal, and a hierarchical Bayesian model for AI statistical inference. In simulation, *airpart* accurately detected partitions of cell types by their AI and had lower RMSE of allelic ratio estimates than existing methods. In real data, *airpart* identified differential AI patterns across cell states and could be used to define trends of AI signal over spatial or time axes.

**Availability:** The *airpart* package is available as an R/Bioconductor package at https://bioconductor.org/packages/airpart.

## 1 Introduction

Measurement of allelic expression (AE) through RNA-sequencing experiments can be used to detect genes for which genetic variation in local *cis*-regulatory elements (CRE) affects cell, tissue, and organism development. Allelic imbalance (AI), in which one allele is expressed higher or lower than the other, may indicate local CRE where regulatory function, e.g. binding of a transcription factor to its motif, is impacted by genetic variation, or could reflect allelic differences in epigenetic state, as in the case of imprinting where maternal or paternal inheritance determines which allele is expressed higher. When allelic expression is quantified in bulk tissue or in a manner lacking the necessary time or spatial resolution, cell-type-specific (CTS) or contextual AI signals may be weakened. As the catalog of accessible CRE and active transcription factors differs across cell lineage, developmental time, and spatial location (Heinz *et al*., 2015), single-cell, spatial, and time-series transcriptomic datasets can help to reveal the cell state or spatial dependencies of genetic effects (Andergassen *et al*., 2017; Wills *et al*., 2013; Combs and Fraser, 2018). For example, it has been observed that allele balance changes dynamically along embryo development stage (Larsson *et al*., 2019; Deng *et al*., 2014) and at HLA genes and other autoimmune loci (Gutierrez-Arcelus *et al*., 2020).

AE analysis cannot detect all variants detectable from expression quantitative trait loci (eQTL) analysis, which examines the association of total expression with genotype, as AE analysis is restricted to those genes and individuals that harbor heterozygous exonic variants (Khansefid *et al*., 2018). However, as total expression level can be affected by technical artifacts (batch effects), environmental effects, or distal-regulation, the within-individual comparisons in AE analysis offers an advantage in focusing on *cis*-regulatory effects, and may increase power (Vigorito *et al*., 2021). Single-cell studies offer a unique opportunity to detect extra cis-eQTLs that would not have be identified in bulk, and hence a number of single-cell studies have been proposed to identify CTS cis-eQTL (Van Der Wijst *et al*., 2018; Cuomo *et al*., 2021b,a). Perhaps due to difficulties in obtaining single-cell AE with sufficient coverage, less attention has been paid to single-cell AE analysis, though single-cell AE analysis can be performed even within a single sample, while single-cell eQTL requires a population of cells of different genotype. Therefore, we get much cleaner signal without dealing with biology or technical noise and a more powerful study design through AE analysis. Recent Smart-seq2 and Smart-seq3 experiments enable full length transcript coverage from single cells at sufficient unique molecular depth to characterize AE for over 10,000 genes. When applied to cells of F1 offspring from crosses of different strains or species, AE data can be generated across hundreds or thousands of cells. (Picelli *et al*., 2014; Larsson *et al*., 2019; Hagemann-Jensen *et al*., 2020)

Prior studies in single-cell AE have categorized genes by allelic state (e.g. bi- or mono-allelic to one or the other allele), estimated allele-specific burst kinetics (Jiang *et al*., 2017), and resolved multimapping reads to genes and alleles in order to reduce spurious monoallelic signal (Choi *et al*., 2019). In this work, we do not address the problem of accurate estimation of AE, assuming access to long reads uniquely assigned to alleles as obtained with Smart-seq or similar technologies (Tian *et al*., 2020). Previous methods have been proposed to detect imprinted genes from single-cell AE (Santoni *et al*., 2017), whereas we focus on detection of CTS *cis* genetic regulation resulting in a consistent imbalance within a group of cells toward a particular allele regardless of parent-of-origin. Furthermore, a recent regression-based method has been proposed to leverage datasets with bulk AE and single-cell total expression of the same tissue, to infer cell-type-specific AI (Fan *et al*., 2021). Here we examine single-cell AE datasets, as well as other contextually-resolved AE data, including spatial or time-series AE. When the AI only exists in one or more specific cell type(s) or the AI varies among cell types, we refer to this phenomenon as differential allelic imbalance (DAI). As single-cell studies providing sufficient coverage and read length for allelic quantification are only now emerging, we are aware of only one related statistical method for detecting DAI, *scDALI* (Heinen *et al*., 2021), which models allele-specific chromatin accessibility using Gaussian Process regression.

Here we introduce *airpart*, an Allelic Imbalance R package for PARTitioning groups of cells, leveraging methods for the Generalized Fhssed Lasso (Devriendt *et al*., 2021) and hierarchical Bayesian modeling, to identify DAI across groups of cells or samples. Our AI models are flexible in terms of the experimental design, and can be applied to group cells or samples by cell type, spatial location or time points, as well as allowing adjustment for covariates. The gene clustering and partitioning of cell types by similar AI signal increases accuracy in the subsequent AE estimation step, alleviating issues from low counts and small numbers of cells for certain cell types or cell states. Our method helps to find subsets of genes sharing similar DAI signals, and helps to generate hypotheses of CTS *cis*-regulatory mechanisms, which can be further validated through experimentation assaying CRE activity or accessibility in particular cell types. The method is available as an R/Bioconductor (Huber *et al*., 2015) package with an accompanying software vignette at https://bioconductor.org/packages/airpart.

## 2 Methods

A summary of the *airpart* workflow is shown in Figure 1. *airpart* takes as input two count matrices and a categorical variable: 1) the allelic counts for the alternate (a1) and reference (a2) alleles across genes (rows) and cells/samples (columns) and 2) the annotated cell types (or spatial location or time point for bulk RNA-seq) in the same order as cells/samples in count matrices. Annotation of cell type can either be provided as prior information or generated by clustering cells by total count (all scRNA-seq experiments analyzed here had prior annotation linking cells to their cell type). The allelic counts could be generated using scBASE (Choi *et al*., 2019), or the quantification pipeline outlined in Larsson *et al*. (2019) for example, for well-characterized diploid transcriptomes. Those inputs are used to construct a *SummarizedExperiment* (Lawrence *et al*., 2013), and functions are provided to determine genes and cells passing quality control (QC). We define the observed allelic ratio as the count ratio of alternate allele reads to the total reads. *airpart* clusters genes with similar AI pattern across cells (see Supplementary Methods Section 1.1 for details). Clustering provides two benefits: it stabilizes DAI detection and estimation in the case that similar patterns occur across genes (e.g. genes under similar patterns of CTS *cis*-genetic regulation), and it speeds up computational time by fitting a partition model (described below) per cluster instead of per gene. In the following methods, we consider one gene cluster at a time. For each gene cluster, a Generalized Fused Lasso framework (Devriendt *et al*., 2021) with Binomial likelihood is used to partition cell types, or a non-parametric method is used. Each of these rely on a graph **Γ** where vertices represent cell types and an edge indicates a pair of cell types that can be fused. *airpart* does not further partition cells within cell types, these are taken as fixed input of the method. DAI is declared if the partition has more than one group of cell types. Given the partition, a hierarchical Bayesian model is fit, which provides allelic ratio estimates and AI statistical inference. *airpart* also includes a number of visualization functions for exploratory data analysis, presentation of partitions and statistical inference on allelic ratios. A summary of the notation used in the following section is provided in Supplementary Table S1.

**Figure 1:**
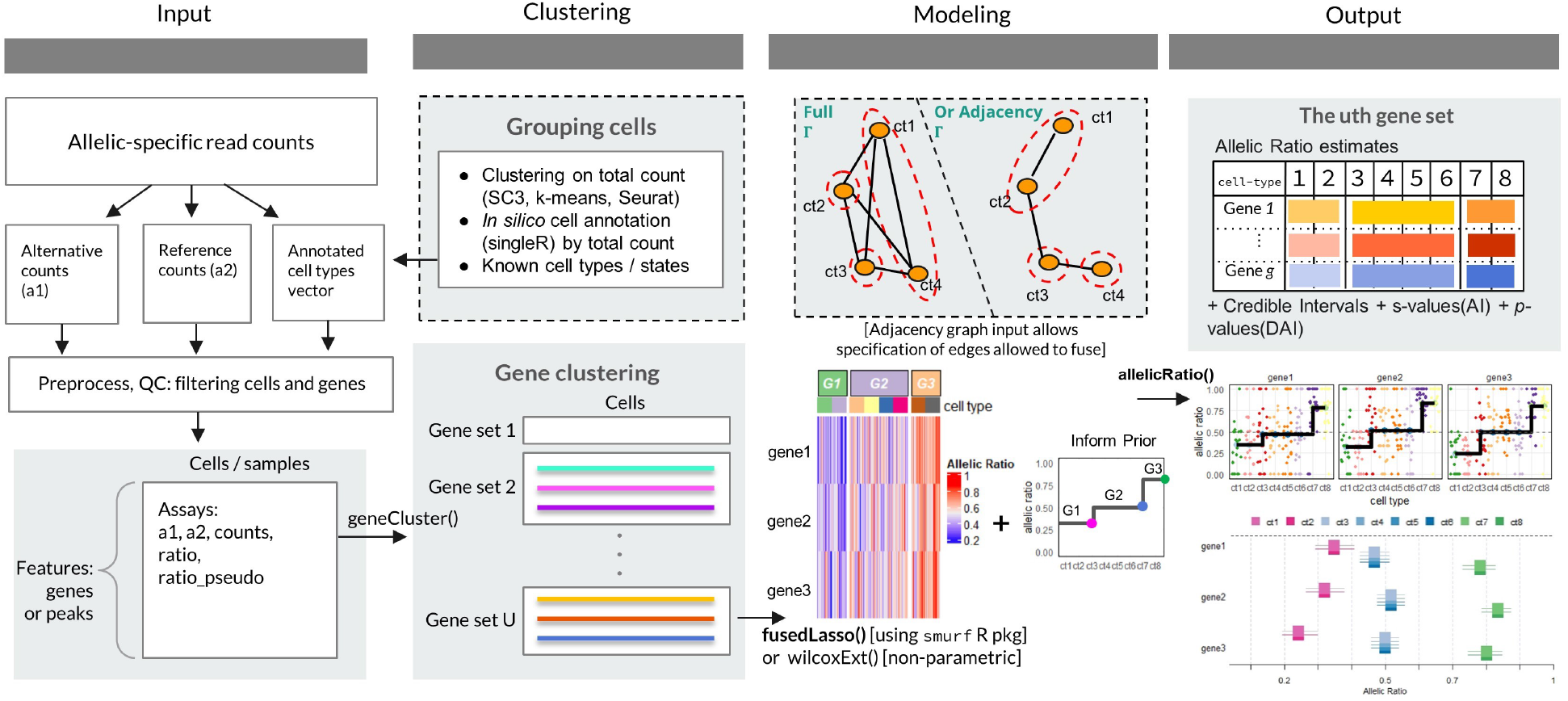
Overview of *airpart* framework. *airpart* takes as input allele-specific read counts, quantified upstream of our method. Known cell annotation or cell clusters derived from total counts are also part of the input to *airpart*. Following QC steps, clustering is performed on genes based on their allelic signal over cells. Then during the modeling step, a partition of the cell groups is generated as shown in heatmap, either by a Generalized Fused Lasso or a nonparametric method. Estimated coefficients of this gene cluster using GFL inform the prior of hierarchical Bayesian model. Finally, *airpart* outputs estimates of allelic ratio for each gene and cell group, as well as s-value or adjusted *p*-value for AI and DAI test, respectively. Multiple visualizations of input data, gene clustering, and fitted parameters are available as functions within *airpart* software.

### 2.1 Distributional assumptions for allelic counts

In previous work, researchers often used a Binomial model (Castel *et al*., 2020) or a Beta-Binomial model for the allelic counts (Skelly *et al*., 2011; Castel *et al*., 2015; Edsgärd *et al*., 2016; Santoni *et al*., 2017; Heinen *et al*., 2021; Choi *et al*., 2019; Zitovsky and Love, 2020), whereas BSCET uses a linear regression for the CTS AI test (Fan *et al*., 2021). For the datasets examined in the Results, either SMART-seq2 single-cell datasets, or spatially- or time-resolved bulk RNA-seq, we found that a Binomial assumption was sufficient for grouping cell types or conditions by AI, as many genes had minimal over-dispersion relative to a Binomial model. However, one real dataset examined in Results exhibited over-dispersion of allelic counts relative to a Binomial, and so non-parametric methods were considered for the partition. When deriving per-gene allelic ratio estimates within a cluster, we modeled allelic counts using a Beta-Binomial generalized linear model. In summary, *airpart* offers Binomial or non-parametric models for partitioning cell types, and Beta-Binomial for deriving allelic ratio estimates.

### 2.2 Generalized Fused Lasso with Binomial likelihood

*airpart* leverages a generalized fussed lasso (GFL) framework (Devriendt *et al*., 2021) implemented in an R package *smurf*, for partitioning cell types into groups of similar allelic ratio. For gene cluster *u*, suppose there are *G* genes, *I* cells and *K* cell types. For gene *g* and cell *i*, let *Y_gi_* indicate the observed allelic count for the alternative allele, *m_gi_* indicate total count, and *r_gi_* = *Y_gi_*/*m_gi_* indicate the observed allelic ratio, *x_i_* indicate cell or sample state which could be cell types, or discrete spatial or time points, and *C_i_* indicate any additional covariates that may associate with allelic ratios. We note that *x* and *c* are represented internally with dummy variables. We assume the following distribution for the alternative allele count with the logit link function

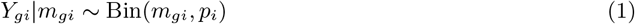

where *p_i_* indicates the true allelic ratio for cell *i*. We define 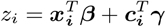, such that

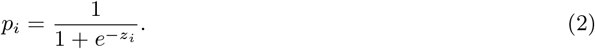

Without loss of generality, we will refer to the values of *x_i_* as cell types. The GFL in *smurf* was used to fit coefficients representing cell-type-specific allelic ratios, where fusing two coefficients means the two cell types are predicted to have the same allelic ratio. Given the graph **Γ** as shown in Figure 1 in the Modeling panel, a complete graph (the default) can fuse all pairwise cell types differences, or alternatively, an adjacency graph can be used with specific edges denoting the cell types that can be fused. The specific edges of **Γ** would be provided *a priori*, for example allowing fusing only among cell types on the same major branch of a developmental trajectory. Suppose *h*, *k* ∈ {1,…, *K*} are any two cell type vertexes that are connected in the graph, then the regularized objective function for model is

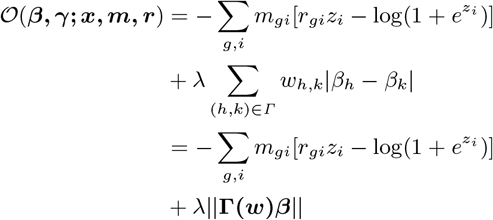

where the second part is the regularization term of the GFL (Höfling *et al*., 2010). Here *β_h_* and *β_k_* denote elements of full parameter vector *β* such that (*β*_0_, *β*_1_,…, *β_K_*) = *β* and **Γ**(*w*) is the matrix with dimensions *N*_Γ_ × *K* where *N*_Γ_ is the total number of edges in the graph **Γ**.

Estimation of coefficients relies on selection of optimal λ and specification of adaptive penalty weights for asymptotic consistency (Devriendt *et al*., 2021). The standardized adaptive weights in *smurf* are defined as 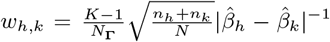 to adjust for possible level imbalances where *n_k_* represent number of cells in cell state *k*. λ was chosen according to the criterion of the lowest deviance (negative of log likelihood) within one standard error of the minimum deviance observed across a grid of λ values, based on 5-fold cross-validation, which encourages parsimony (more fusing of pairwise differences). When there are less than 8 cell types represented in *x*, λ was chosen by finding the lowest deviance within half a standard error of the minimum, thus allowing more non-zero pairwise differences to persist. In addition to the Binomial likelihood(*airpart.bin*), a Gaussian likelihood(*airpart.gau*) was considered, which assumes *r_gi_* ~ *N*(*p_i_*,*σ*). The different likelihood models for GFL were compared via simulation.

### 2.3 Pairwise Mann-Whitney-Wilcoxon Test

An alternative method is considered and available within *airpart*, relying on a nonparametric test(*airpart.np*), both for increased speed and for cases when the distributional assumptions of the above model do not fit the data. We extend the Mann-Whitney-Wilcoxon (MWW) test to derive a partition based on the p-values from pairwise comparisons across cell types.

For all pairs of cell types with edges in **Γ**, pairwise MWW tests were performed for the allelic ratio distribution difference. A similarity score matrix 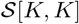 was constructed, with elements equal to the MWW test p-values. Each element of this matrix was therefore related to the separability of the ranked allelic ratios for the two cell types. This matrix was then binarized into 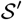 as follows: 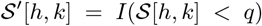 with *I*() the indicator function. This binarization depended on a tuning parameter *q* and defined a network adjacency matrix. For pairs not represented by edges in 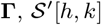 was set to 0. Finally the adjacency matrix was used as input as a distance matrix for hierarchical clustering.

To choose the tuning parameter *q* and find the cell types partition, a model selection was performed based on the Bayesian Information Criterion (BIC). The BIC scores a candidate model using both its performance on the in-sample error and the complexity of the model. The best model was chosen by minimizing a loss function defined below, along a range of *q* = 10^*v*^ where *v* = −2, −1.8,…, −0.4. The loss function was constructed based on the Gaussian special case of BIC that assumes independent errors from a normal distribution, and that the derivative of the log likelihood with respect to the true variance is zero (Hannan, 1982). We have

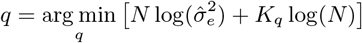

where N is the total number of elements within this gene cluster (*N* = *G* × *I*), 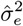 is an estimate of the error variance, and *K_q_* is the number of groups derived from constructing adjacency matrix according to each *q* threshold among the partition of *K* cell types. The estimate of the error variance in this case is defined as 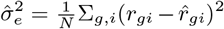, which is a biased estimator for the true variance. In terms of partition group the loss function is

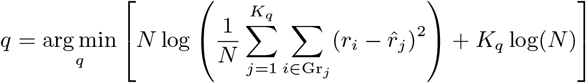

where Gr_*j*_ is a set of cells in group *j*, 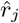 is the mean allelic ratio of all elements within group *j*.

### 2.4 Hierarchical Bayesian modeling

In the *airpart* steps to estimate the partition of cell types within a gene cluster, the true allelic ratio *p_i_* being modeled with the GFL did not vary across genes within the cluster. However, the allelic ratio may vary across genes within a cluster, though the clustering step brings together genes with similar patterns of allelic ratio. To derive per-gene allelic ratio estimates and credible intervals within a gene cluster, a Bayesian Beta-Binomial generalized linear model (GLM) was used. This GLM can be fit sharing prior information for all cells within a cell-type group defined by the partition (“grouped”) or one cell type at a time, ignoring the partition (“nogroup”). The case of ignoring the partition allows for estimation of allelic ratios even if the input *x* is continuous-valued. Let *ϕ_g_* indicate a gene-specific dispersion parameter, then we assume the following model:

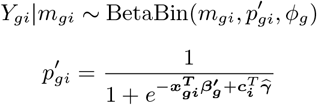

where 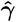 is the *maximum a posteriori* (MAP) estimate derived from *apeglm* (Zhu *et al*., 2018; Zitovsky and Love, 2020) and used as offset in the model (if covariates are provided). The dispersion *ϕ_g_* controlling the variance of *Y_gi_* by:

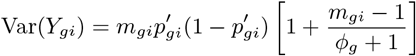

Shrinkage estimation was performed separately on the dispersion parameter and coefficients representing cell type allelic ratios. We first describe shrinkage on the dispersion parameter. We assume the logarithm of the dispersion estimate 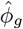 follows a Normal distribution,

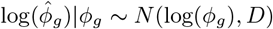

where D, the sampling variance, is assumed equal across all genes in the cluster. We estimated this sampling variance with 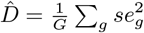 where *se_g_* is the estimated standard error of the logarithm of the dispersion estimate 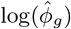. Both 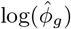 and *se_g_* were estimated using the *apeglm* (Zhu *et al*., 2018) software. Although the dispersion parameter was estimated per gene, the gene-wise models were linked by global hyper-parameters which were estimated from the entire gene cluster at once. The specification of a cluster-specific prior was used a simple means of sharing information between genes. We assume that the dispersion parameters log(*ϕ_g_*) follow a Normal distribution

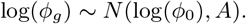

log(*ϕ*_0_) was estimated from the gene-wise MLEs, 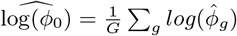, and the variance was estimated with 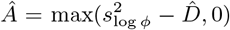 where 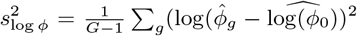. In order to obtain the relative weighting of the gene-wise and global variance estimators, 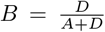 was defined as a parameter to shrink dispersion estimates towards a middle value (roughly following Efron and Morris (1975)). The final estimate for dispersion used in fitting posterior coefficients is:

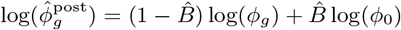

where 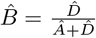.

A Cauchy distribution was used as the prior for *β_g_* the coefficients representing cell type allelic ratios. Shrinkage estimation was performed one group at a time, where a group is defined by the cell types within a partition from the first step or alternatively, ignoring groupings, meaning each cell type is estimated by itself. Without loss of generality, we describe the grouped case. Let *μ* define the center of the prior distribution for *β_g_*, which will be a vector of *J* if there are *J* cell type groups in this gene cluster. The GFL estimates 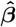 were used as 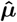 across the multiple genes within a cluster (or weighted means were used if nonparametric methods were used for defining the partition in the previous step). For the estimation of per gene ratios, we assume the coefficients follow a Cauchy distribution,

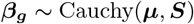

where *S* is scaling parameter estimated as part of the *apeglm* method (Zhu *et al*., 2018), and 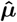 was plugged in as the center of the prior distribution. Maximum posterior estimates and credible intervals are estimated for a Beta-Binomial likelihood using *apeglm*. To assess allelic imbalance across each cell type and each gene within a cluster, *s*-values (Stephens, 2017) were calculated and provided, where thresholding on this value provides control of the aggregate false sign rate (the rate of incorrect signs of estimates within the reported set). We note that while the credible intervals and *s*-values calculated in this step reflect uncertainty in estimation of the allelic ratio based on number of cells and the range of the counts, as we fix the gene clustering and cell type partition from previous steps, uncertainty from those upstream steps is not included in the inference provided by the hierarchical model.

### 2.5 Simulation setup

In order to assess *airpart*’s partitioning of cell types by allelic ratio, and its accuracy of estimates of the allelic ratio itself, we performed three sets of simulation tests. The allelic counts were simulated from a Beta-Binomial (BB) distribution with constant dispersion parameter *ϕ* for all genes, so we ignore the index *g* here. The total counts were drawn from a Negative Binomial (NB) distribution. Half of the total counts had a mean count of 2 while half of the total counts had a higher mean count, ranging across different simulations among values of *cnt* ∈ {5, 10, 20}. In each case, the NB dispersion was set to *α* = 5 (dispersion α defined such that Var(*Y*) = *μ* + *αμ*^2^). As *airpart* combines allelic counts from multiple genes when finding the partition of cell types, having lower and higher total counts for each gene is equivalent to a dataset with a mix of low and high count genes within a gene cluster. The mean counts and BB dispersion values (*ϕ* = 20) were chosen based on estimated parameters over real SMART-seq2 scRNA-seq datasets (Larsson *et al*., 2019), as shown in Figure S1(A). However, we observed that in earlier datasets, such as Deng *et al*. (2014), as shown in Figure S1(B), the allelic counts generated lower *ϕ* estimates (more variance). Thus we also assessed simulations using *ϕ* = 3 to evaluate method robustness when the data was substantially overdispersed relative to a Binomial model (the model used by the GFL in *airpart*). *airpart* was assessed via simulation across various settings summarized in Supplementary Table S2, and compared to another statistical method for detecting heterogeneity of allelic ratio in scRNA-seq, *scDALI* (Heinen *et al*., 2021), which models allele-specific chromatin accessibility using Gaussian Process regression.

The adjusted Rand index (ARI) was used to assess accuracy of the cell type partitions with respect to the true partition by allelic ratio. The number of gene within a gene cluster was varied across *g* ∈ {5, 10, 20}. *ARI* ∈ [−1, 1] measures the corrected-for-chance version of the correct partition of the cell types. ARI of 1 means a perfect partition, and an ARI of 0 is no better than random guessing. Each cell type was simulated to have 40 cells, and 10 cell types were simulated (400 cells in total) with true allelic ratio given by {0.95, 0.9, 0.85, 0.85, 0.7, 0.7, 0.65, 0.6, 0.5, 0.5}, thus with a true partition of seven groups of unique allelic ratios. The whole simulation was repeated 200 times for each combination of simulation parameters: high total count (*cnt*), number of genes (*n*), and dispersion value (*ϕ*).

For evaluating allelic ratio estimation accuracy, 400 genes were simulated such that all have the same pattern of DAI. The number of cells per cell type was varied from 40 to 100. RMSE was used to evaluate performance. Here, we additionally assessed the effect of using the partition to aid in estimation accuracy, by comparing performance with and without this cell type grouping step. The version of the estimation method without the grouping step was denoted as *airpart.nogroup*. We define a simulation parameter *d* ∈ {0.05, 0.1, 0.2, 0.3}, and at each *d*, allelic ratios according to {0.3, 0.3, 0.3 + *d*, 0.3 + *d*, 0.3 + 2*d*, 0.3 + 2*d*, 0.3 + *d*, 0.3 + *d*} were simulated following a U-shaped pattern, where *d* controls the extent of the rise and fall in allelic ratio. For this simulation, the cell type indicator *x* was provided as a matrix with one-hot encoding to *airpart* and *scDALI*. We used *scDALI* with a linear kernel which assumes that the allelic ratio has a linear trend across covariates. which in this case allows for estimation of CTS allelic ratios.

For significance testing of DAI, genes were simulated without DAI {0.5, 0.5, 0.5, 0.5, 0.5, 0.5} and with DAI {0.5, 0.5, 0.6, 0.6, 0.7, 0.7} on 6 cell types with 40 and 100 cells per cell type, separately. For *airpart*, a likelihood ratio test was performed to compare the model with multiple groups and to a reduced model with an intercept only. We ran *scDALI* using `scDALI-Het` which calculated score test statistics. We adjusted allelic heterogeneity p-values for both methods using the Benjamini-Hochberg (BH) (Benjamini and Hochberg, 1995) procedure with a cutoff of 0.05.

### 2.6 Allelic datasets

We applied *airpart* to 2 single-cell RNA-seq datasets: Larsson *et al*. (2019); Deng *et al*. (2014) and 2 bulk RNA-seq datasets (Gutierrez-Arcelus *et al*. (2020); Combs and Fraser (2018)). From those datasets, Larsson *et al*. (2019) contains 224 mouse embryo stem cells (C57BL/6 × CAST/EiJ) and 188 mouse embryo fibroblasts (CAST/EiJ × C57BL/6J) grouped across states of cell cycle (G1, S, G2M), as identified by the authors. Deng *et al*. (2014) includes 286 pre-implantation mouse embryo cells composed of 10 cell types from an F1 cross of female CAST/EiJ and male C57BL/6J((B6)) mice. Cells were sampled along a time course from the zygote and early 2-cell stages through the late blastocyst stage of development. Maternal allelic ratios were estimated for the two scRNA-seq datasets. Gutierrez-Arcelus *et al*. (2020) stimulated memory CD4+ T cells from 24 genotyped individuals of European ancestry with anti-CD3/CD28 beads and characterized the dynamics of AI events at 0, 2, 4, 8, 12, 24, 48 and 72 hours after stimulation. Combs and Fraser (2018) performed RNA-seq of five hybrid *Drosophila* species *D. melanogaster* × *D. simulans* embryos sliced along their anterior-posterior axis to identify genes with spatially varying allelic imbalance. Results applying *airpart* to Combs and Fraser (2018) dataset are provided as Supplementary Results. The cell population annotations for all datasets were provided with the data. These annotations were used as known cell types/states/spatial position for analysis. The number of cells in Table 1 represents the size of each dataset after preprocessing (see Section 1.2 for details).

**Table 1:** Single-cell (sc) and bulk RNA-seq datasets used for evaluation.

| Source | Observations | States/<br>Contexts | Tissue type |
| --- | --- | --- | --- |
| Larsson *et al.* (sc) | 367 | 4 | Mouse F1 embryos |
| Deng *et al.* (sc) | 228 | 10 | Mouse F1 embryos |
| Gutierrez-Arcelus *et al.* | 199<sup>1</sup> | 8 | Mem. CD4+ T cells |
| Combs *et al.* | 126<sup>2</sup> | 19 | Fly F1 embryos |

<sup>1</sup> time point replicates from 24 donors;
<sup>2</sup> slices from 5 embryos.

### 2.7 Functional enrichment analyses

In order to check whether gene clusters with specific differential allelic imbalance pattern detected by *airpart* were enriched for functional categories or were correlated with enhancer activity, we performed GO term analysis and genome enrichment analysis. GO term enrichment was calculated using the *goseq* package (Young *et al*., 2010) with the UCSC mm9 gene lengths database. Along with the single-cell dataset of (Larsson *et al*., 2019), there was additional chromatin immunoprecipitation and sequencing (ChIP-seq) data for H3K27ac for one of the parental lines (B6) for mESC and fibroblasts. ChIP-seq peaks were selected with fold enrichment > 15. As ChIP-seq data was only available for the B6 strain, we assessed whether the genes with allelic imbalance toward one allele were more closely associated with enhancer activity in that cell type compared to the other cell type by Fisher’s exact test.

## 3 Results

### 3.1 Simulation

We evaluated *airpart* across a variety of simulation datasets, and in comparison to a newly developed method for detecting heterogeneous allelic imbalance, *scDALI* (Heinen *et al*., 2021). On the simulated dataset with 10 cell types of different allelic ratio, as described in Section 2.5, the Generalized Fused Lasso (GFL) with Binomial likelihood tended to have higher ARI than the GFL with Gaussian likelihood or our nonparametric method when the number of genes within a gene cluster was less than 20 (Figure 2A). The higher ARI here indicates that a method is more accurate at partitioning the cell types according to their true underlying allelic ratio. This result aligned with that of previous work showing that modeling the allelic ratio using count distribution can increase power (Sun, 2012). When the total number of observations (20 genes × 40 cells per gene) was greater than 800, both GFL with Binomial likelihood and the nonparametric method almost always achieved ARI of 1 regardless of the value of higher total count (*cnt*). In such cases, running the nonparametric method is preferred as the computation time was around 6 time faster than `airpart.bin` (Figure S2A). The nonparametric method had higher ARI and shorter whisker in the case of more overdispersion relative to a Binomial (*ϕ* = 3) which was expected (Figure S2B).

**Figure 2:**
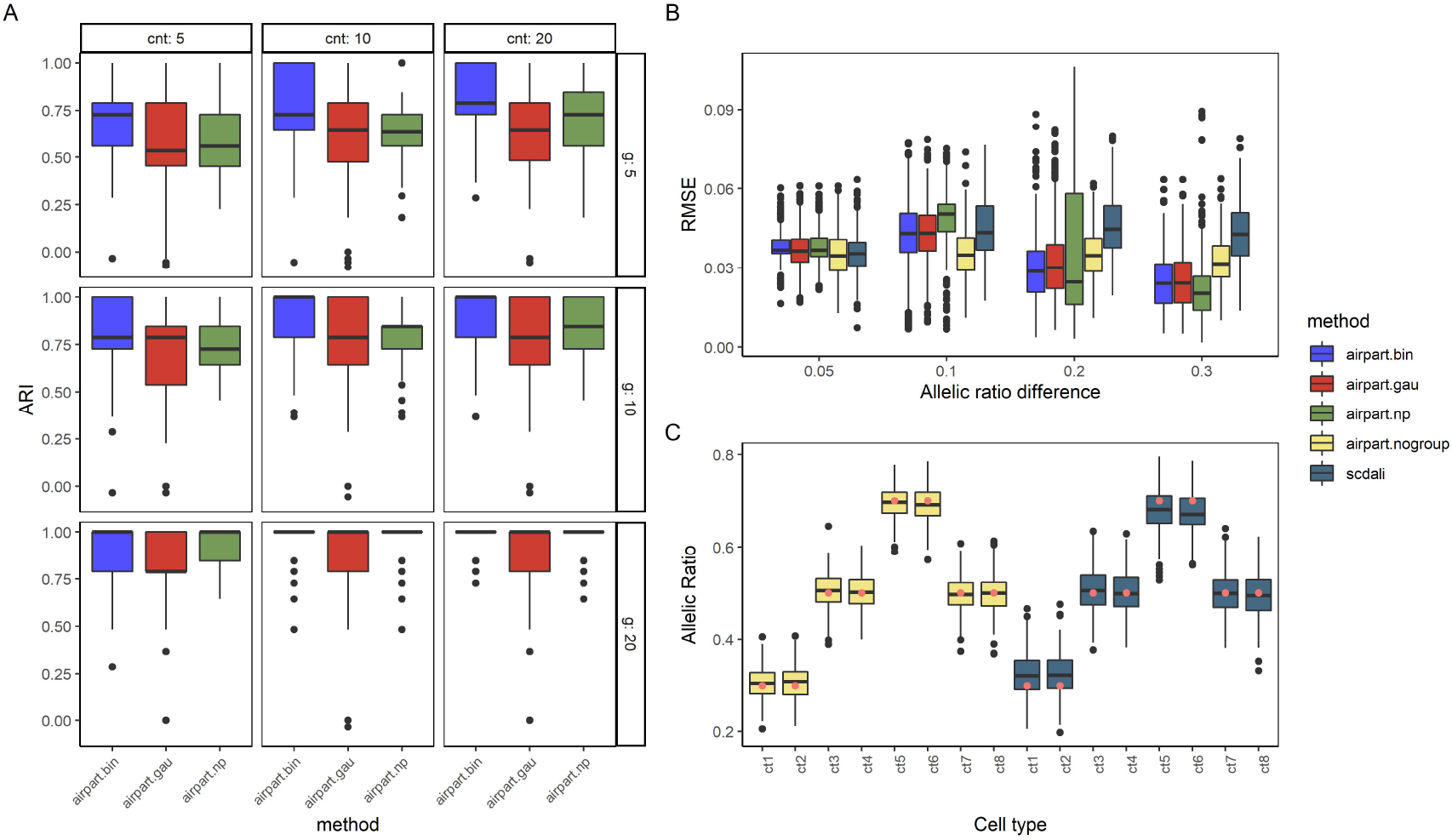
Performance comparison of *airpart* variants and *scDALI* on simulation datasets. A) Boxplot of partition accuracy among 3 variants of *airpart*. y axis is adjusted Rand index among 200 iterations. cnt: the higher mean count, n: number of genes within a gene cluster. B) Boxplot of RMSE per gene for estimation of the allelic ratio for *n* = 40 cells among 400 iterations. Each gene has an underlying U-shape pattern described in the Section 2.5. C) Boxplot demonstrating *airpart* without cell type grouping step and *scDALI* performance on each cell type at DAI = 0.2. The red dots represent the true allelic ratio values of {0.3, 0.3, 0.5, 0.5, 0.7, 0.7, 0.5, 0.5}.

The simulated total counts distribution mimicked real scRNA-seq counts distribution (Figure S1C), and *airpart* clustered together genes for which the allelic ratio trend was similar (Figure S1D).

We compared the allelic ratio estimation accuracy of *airpart* with or without grouping step to *scDALI* across different scales of DAI. *airpart.bin* had smaller RMSE than *scDALI* for allelic difference > 0.2 while performing comparably for allelic difference of 0.1, and slightly worse for allelic difference of 0.05 (Figure 2B). In this simulation, we assessed one gene each time, so *airpart* did not benefit from aggregating signal across multiple genes. All three variants of *airpart* with partition step (Binomial likelihood, Gaussian likelihood, and non-parametric approach) showed distinct decreasing RMSE with increasing allelic ratio difference, while *airpart.nogroup* (without cell type partitioning) and *scDALI* had relatively constant RMSE. *airpart* variants with partition step benefited in this simulation from its approach toward discrete groupings of cell types, as the simulated data consisted of eight cell types falling in three groups by their true allelic ratio; as the allelic ratio increased, the correct partition was easier to identify, which led to the decrease in RMSE for those three method variants. In order to understand why *scDALI* tended to have slightly larger RMSE than *airpart.nogroup*, we examined the estimates themselves over the cell type variable (*x*); *scDALI*’s estimates tended to shrink towards 0.5 at the extremes on this simulation (Figure 2C). From this simulation, we inferred that when the true model is one of discrete allelic ratios shared across a partition of the cell types, grouping cell types with similar allelic ratio increases observation size and may therefore aid in estimation.

We performed simulation with more cells per cell type (*n* = 100 compared to *n* = 40 in previous simulations) to confirm that estimates would have reduced error with more observations. Both *airpart* and *scDALI* had lower RMSE when *n* = 100 (Figure S3A). *airpart.bin* additionally had better performance relative to *scDALI* for DAI = 0.1, compared to the *n* = 40 simulation. When considering credible interval coverage, *airpart.nogroup* had the highest empirical coverage (the average number of times credible intervals contained the true value) almost always achieving 95%, although other *airpart* variants and *scDALI* performed as well when DAI > 0.1 (Figure S3B,C). Again note that *airpart* partitioned cell type group per gene under this set up, so it didn’t benefit borrowing information from other genes with similar allelic pattern. *scDALI* performed similarly as grouped *airpart* variants when DAI = 0.1.

In the simulation assessing the rate of DAI calling when *n* = 40, *airpart.nogroup* and *scDALI* both had the highest specificity of around 97% compared to other *airpart variants(airpart.bin* had 93.75%). But all methods had similar sensitivity of around 98.00%(Supplementary Table S3). For *n* = 100, *airpart.bin* had the highest specificity of 98.25%, likely due to its increasing accuracy in determining the partition of cell types (Supplementary Table S4). *scDALI* and other variants’ specificity basically didn’t change as *n* increased to 100. But all methods had 100% TPR at this sample size. *aipart.gau* and *airpart.np* had lower specificity in this global test of DAI (89.5% and 88.0% at *n* = 40, and 85.25% and 88.25% at *n* = 100), which was expected as these variants often detected too many groups in the partition analysis, with lower ARI relative to *airpart.bin* (Figure 2A). We expect *airpart.np* would perform better relative to other methods when the data is highly over-dispersed (Figure S2B). Overall, *airpart.bin, airpart.nogroup*, and *scDALI* recovered most DAI while not falsely calling too many genes as DAI.

Overall, we recognize that *airpart* and *scDALI* have subtly different inference goals, with *airpart* predominantly focused on characterizing the allelic ratio patterns that result from discrete groups of cell types sharing a common regulatory context (for example, expression of transcription factors and accessibility of CRE). On the other hand *scDALI* is more suitable for detecting various types of heterogeneous AI including continuous gradients of AI in cells across measured or inferred dimensions.

### 3.2 Mouse ES cells and fibroblasts

For assessing *airpart* on real allelic datasets, we first examined two single-cell RNA-seq datasets consisting of mouse embryo cells (Larsson *et al*., 2019; Deng *et al*., 2014). Both datasets were mouse F1 nonreciprocal crosses in which we observed clusters with allelic imbalance towards the maternal allele, likely from imprinting in mature cells or genome activation for early cell stages. A complete graph was applied to both datasets, allowing any developmental time point coefficients to be fused with another. In both cases, *airpart* partitioned the cell stages as expected according to developmental time, e.g. consecutive and related time periods being fused together, such as early, mid, and late-blastocyst.

*airpart* was first applied to the Larsson *et al*. (2019) dataset consisting of four cell states including three cell cycle states of primary mouse fibroblast (G1, S, G2M) and mouse embryonic stem cells (mESC). After QC filtering, 2,481 genes remained and five gene clusters were detected. Four of the five clusters, comprising 412 genes in total, showed evidence of DAI by their *airpart* partitions (Figure S4A). One cluster of 128 genes partitioned the cell states such that all cell cycles of fibroblast were grouped together and apart from the mESC (Figure 3A and Figure S4C). In this cluster, the fibroblast group had mean estimated allelic ratio around 0.45 and the mESC had allelic imbalance with a ratio of around 0.70 toward the maternal allele. We estimated the allelic imbalance in both mESC and fibroblast and calculated 95% credible intervals (Figure 3B). With an s-value threshold of 0.005 (Section 2.4), all 128 genes demonstrated AI in mESCs and 85 genes out of 128 demonstrated AI in fibroblasts, which was roughly consistent with credible intervals not overlapping an allelic ratio of 0.5. To check whether this cluster had any functional association with stem cell maintenance, gene set enrichment analysis was performed. The most significant Gene Ontology (GO) term was “response to leukemia inhibitory factor (LIF)” (GO:1990823, odds ratio = 4.90, *adj. p* = 0.136 (BH)), where *Lif* is a cytokine involved in embryonic stem cell self-renewal (Hirai *et al*., 2011).

**Figure 3:**
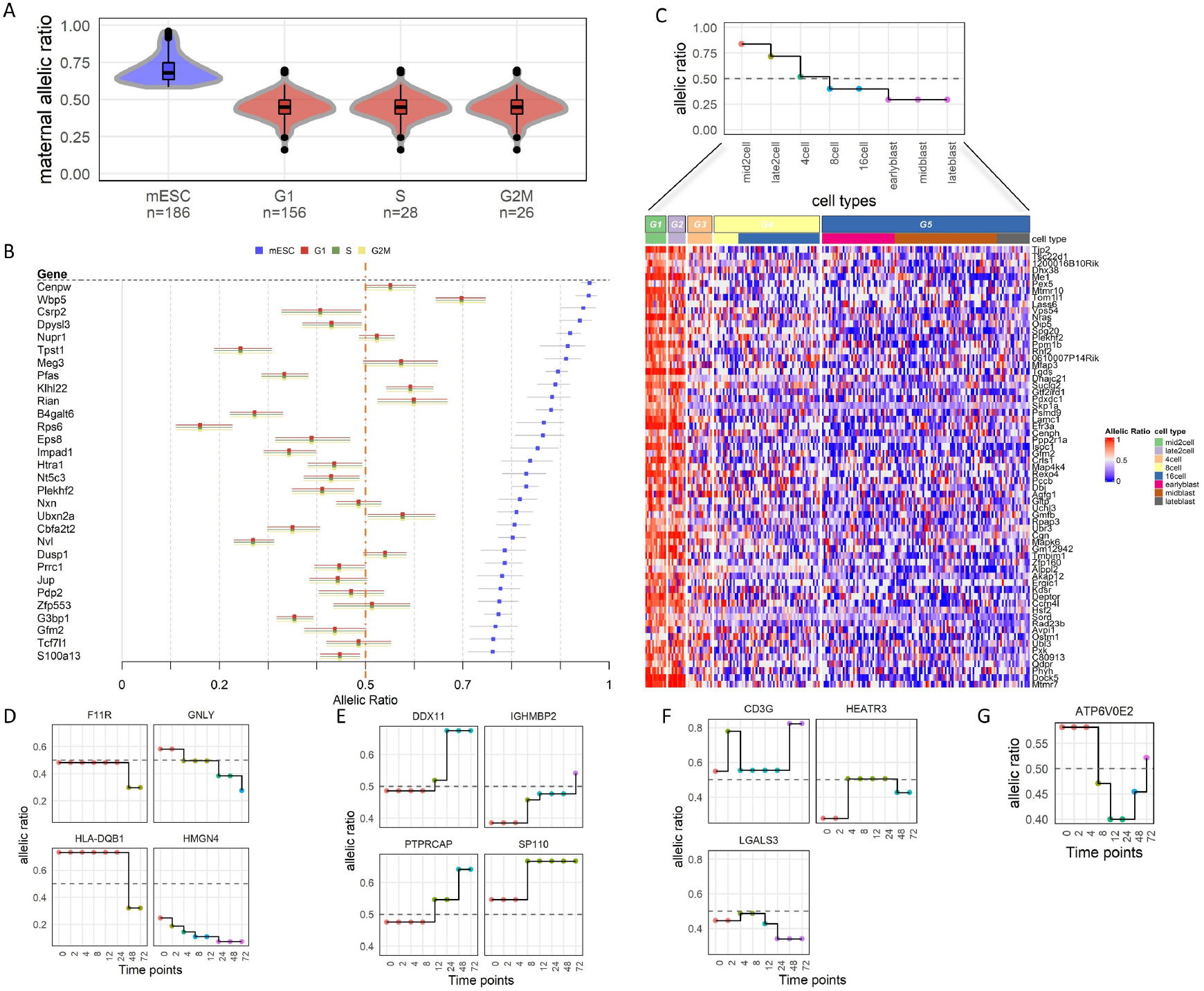
Evaluation of airpart on scRNA-seq and time course RNA-seq experiments. A) Violin plot of estimated allelic ratio on Larsson’s dataset with *n* indicating the number of cells. Color represents different partition groups. B) Forest plot for Larsson’s dataset, showing top 40 genes with smallest *s*-value. Dotted line denotes allelic ratio = 0.5 C) Step plot and heatmap of results for Deng’s dataset. This gene cluster partitioned into 5 groups denoted by color in the step plot. D-G) Selected genes displaying *airpart* fitted model on Gutierrez-Arcelus’s data: (D) decreasing trend, (E) increasing trend, (F) up-down pattern, and (G) down-up pattern.

To assess whether the 128 gene cluster with DAI had an association with enhancer activity, an enrichment analysis was performed using H3K27ac peaks exclusively measured in the B6 strain in both cell types. While the enhancer activity signal was therefore not an allelic signal, we hypothesized that genes which favor the B6 (maternal) allele in mESC may tend to overlap with H3K27ac in B6 in mESC. For the 128 gene cluster with DAI toward maternal allele in mESC, we found that genes in the cluster were more often associated with enhancer activity in mESC compared to fibroblast (significance assessed by Fisher’s exact test *p* = 0.0002).

*airpart* was additionally applied to the Deng *et al*. (2014) dataset consisting of 228 pre-implantation mouse embryo cells passing our QC steps, from an F1 cross of CAST/EiJ × C57BL/6J mice. Ten cell types were annotated from the zygote and early 2-cell stages through the late blastocyst stage of development. Allelic ratio was defined as maternal (CAST / (B6 + CAST)) and 4,679 genes passed our gene QC steps. All genes revealed DAI patterns as expected due to genome activation from zygote onward to the blastocyst stages. In order to focus on DAI after zygote or early 2-cell stages, we performed clustering of genes and partitioning of cell stages with those cell stages removed. 13 out of 16 gene clusters, consisting of 3,019 genes in total, showed DAI pattern after removing these cell stages. One gene cluster showed a decreasing allelic ratio pattern along developmental time and *airpart*’s idea was illustrated by estimating each gene’s allelic ratio through using the gene cluster fused lasso coefficients as prior mean (Figure 3C). The corresponding violin plot of allelic ratios is shown in Figure S4C. The estimated allelic ratio around 0.3 in early/mid/late blast stage was an exception (most clusters have the blastocyst cells as balanced allelic expression). This cluster of 65 genes had significant enrichment for GO terms such as “cell development” (GO:0006139, odds ratio = 2.02, *adj. p* = 0.0017 (BH)) and “cell differentiation” (GO:0030154, odds ratio = 1.29, *adj. p* = 0.0233 (BH)).

### 3.3 Dynamic allelic imbalance during T cell activation

We applied *airpart* to an RNA-seq dataset of stimulated memory CD4+ T cells of 8 discrete time points (Gutierrez-Arcelus *et al*., 2020). To do so, we created a graph *Γ* with edges only between consecutive time points, restricting the fusion of coefficients in the Generalized Fused Lasso. From visualization of PCA plots, and allelic ratio heatmaps of prospective clusters, we determined that genes often had distinctive patterns of allelic imbalance after T-cell stimulation; we recommend such exploratory analysis before the modeling step and provide visualization functions in the associated Bioconductor package. We therefore decided to perform partitioning and allelic ratio estimation gene-by-gene. As there were too few observations to robustly select via cross-validated deviance per gene, we manually set the maximum regularization parameter λ and derived the optimal λ controlling the amount of fusing of time points (Section 2.2) according to BIC within specific range. Following the convention used in Gutierrez-Arcelus *et al*. (2020), we define the allelic ratio as the proportion of reference over total read counts for this experiment. To find most potential genes with dynamic allelic imbalance, we calculate the weighted average allelic ratio for each time point(weighting by the cell’s total count) and each gene, then compute the variance over time points. Among the 43 genes with largest variance of that, most of them were enriched within autoimmune loci, as reported by the original study authors. Examples include *F11R*, a ligand for integrin alpha-L/beta-2 involved in memory T-cell and *HLA-DQB1*, a member of the human leukocyte antigen (HLA) complex. As stated in Methods, we ignore the allelic complexity of the HLA genes in this analysis and group the alleles together into two, based upon the SNP with the largest total count. We chose to reduce to diploid allelic counts in each individual based on a single SNP since HLA typing and across-donor inference of more than two alleles is out of the scope of this study. This approach was used for method demonstration only. *airpart* partitioning of the time series by allelic ratio revealed 4 types of patterns (decreasing, increasing, up-peak and down-peak) as shown in Figure 3 (D,E,F,G) respectively (step-plots for all 40 genes provided in Figure S7). As in the original study, we also observed the dominant allele could switch over the time course, or bi-allelically expressed genes could switch to dominant by one or the other allele. While the original paper used logistic regression with polynomial terms for time within each individual, we recovered similar DAI trends for many autoimmune genes, such as *GNLY* and *DDX11*. Overall, *airpart* successfully captured the DAI patterns seen across T cell activation.

In summary, when applied to scRNA-seq and bulk RNA-seq datasets, *airpart* was able to identify relevant partitions of cell types or samples, with gene clusters displaying significant with biologically meaningful gene sets and cell-type-specific enhancer activity. Results applying *airpart* to a dataset of spatial transcriptomic fly cross embryos Combs and Fraser (2018) are provided in Supplementary Section 2.1, where *airpart* was used to analyze genes based on spatially-varying allelic imbalance.

## 4 Discussion

An understanding of how individual genes may be regulated across context or condition helps to elucidate molecular mechanisms underlying complex phenotypes or diseases, and how heritability of risk may be conferred. Context-specific AE enables isolation of *cis*-acting genetic regulation of transcription, and the study of AE is a good complement to differential gene expression studies, where a multitude of factors may influence differences in total expression across condition. Single-cell RNA-seq of F1 crosses enables measurement of context-specific AE, where the cell type or cell stage can be taken as the context that influences *cis*-genetic regulation. Additionally, spatially resolved or time-series allelic datasets offer another such example. Context-specific or conditional allele specific expression datasets can detect allelic imbalance with fewer samples than context-specific quantitative trait locus (QTL) studies (Findley *et al*., 2021), although measurement of allele specific expression in a sample requires presence of heterozygous variation in the transcribed region, which may not occur for all transcripts or for all genes depending on the population under study. With the advent of large-scale systematic assays for interrogating variants and regulatory elements, such as CRISPR-Cas9 and massively parallel reporter assays (MPRA), there are now increasing opportunities to re-use context-specific allelic datasets which can help point to the key cell types or cell states for validation.

To assist with analysis of such datasets, we developed *airpart*, a statistical framework for identifying genes and cell types or cell stages with similar DAI signal. We refer to the partitioning step of *airpart* as an “interpretable” model, in that the discrete grouping of cell types can be interpreted as a hypothesis that they share a common mechanism of *cis*-regulation, such as a common set of expressed transcription factors and accessible set of CRE. As scRNA-seq data often have low counts for some genes of interest, and as the experiments used for allelic expression in single cell (SMART-seq2 or SMART-seq3) often have a relatively small number of cells per cell type, we employed gene clustering and a partitioning of cell types in order to increase power. Aside from using clustering to detect meaningful subsets of genes by allelic imbalance, it is also possible to provide pathways or other gene sets known *a priori* for *airpart* to partition by allelic imbalance signal as a group. In this way, *airpart* stabilizes gene-level estimation by borrowing information about the similarity of cell types from other genes that have similar AE patterns.

*airpart* can be applied to a variety of problems, as it leverages a Generalized Fused Lasso framework (Devriendt *et al*., 2021) where a graph specifying the connectivity of the cell types is provided, wherein only certain cell types can be allowed to fuse, helpful for scenarios such as time-series experiments or for prohibiting fusing across different cell lineages. Another point of flexibility is *airpart*’s use of a design matrix within the generalized linear model (Eq. 2) wherein additional covariates can be provided that may also have effects on the allelic ratio. This was used here in the analysis of the time course RNA-seq dataset to adjust for individual effects, and may be helpful for multi-individual single-cell sequencing studies. Furthermore, *airpart.no group* can accept a design matrix representing natural cubic splines (Hastie, 1992). *airpart* therefore offers fast estimation of smooth functions of the allelic ratio over a continuous variable, making use of a hierarchical model to stabilize the over-dispersion parameter (Supplementary Section 2.1). Though *airpart* predominantly focuses on characterizing the allelic ratio patterns that result from discrete groups of cell types under shared regulatory contexts, *airpart* can in this way be used to model continuous gradients of *cis*-regulatory effects on cells or samples.

## 5 Availability

*airpart* is implemented as an R/Bioconductor package available at: https://bioconductor.org/packages/airpart. The package contains a detailed vignette demonstrating use of the functions. All of the R code and data used in this paper for evaluating methods on simulated and real RNA-seq datasets, as well as HTML vignettes reproducing paper figures for analysis of Larsson *et al*. (2019) and Deng *et al*. (2014), are available at the following repository: https://github.com/Wancen/airpartpaper.

## Acknowledgments

We thank Peter A. Combs and Hunter B. Fraser for providing guidance on processing of the *Drosophila* embryo spatial dataset. We thank Anna Alessandra Monaco for helpful discussion on allelic single-cell datasets.

## References

Andergassen, D., Dotter, C. P., Wenzel, D., Sigl, V., Bammer, P. C., Muckenhuber, M., Mayer, D., Kulinski, T. M., Theussl, H.-C., Penninger, J. M., and et al. (2017). Mapping the mouse Allelome reveals tissue-specific regulation of allelic expression. eLife, 6.

Benjamini, Y. and Hochberg, Y. (1995). Controlling the false discovery rate: a practical and powerful approach to multiple testing. Journal of the Royal statistical society: series B (Methodological), 57(1), 289–300.

Castel, S. E., Levy-Moonshine, A., Mohammadi, P., Banks, E., and Lappalainen, T. (2015). Tools and best practices for data processing in allelic expression analysis. Genome biology, 16(1), 1–12.

Castel, S. E., Aguet, F., Mohammadi, P., Ardlie, K. G., and Lappalainen, T. (2020). A vast resource of allelic expression data spanning human tissues. Genome biology, 21(1), 1–12.

Choi, K., Raghupathy, N., and Churchill, G. A. (2019). A Bayesian mixture model for the analysis of allelic expression in single cells. Nature communications, 10(1), 1–11.

Combs, P. A. and Fraser, H. B. (2018). Spatially varying cis-regulatory divergence in Drosophila embryos elucidates cis-regulatory logic. PLOS Genetics, 14(11), 1–23.

Cuomo, A. S., Heinen, T., Vagiaki, D., Horta, D., Marioni, J., and Stegle, O. (2021a). Cellregmap: A statistical framework for mapping context-specific regulatory variants using scrna-seq. bioRxiv.

Cuomo, A. S., Alvari, G., Azodi, C. B., McCarthy, D. J., Bonder, M. J., et al. (2021b). Optimising expression quantitative trait locus mapping workflows for single-cell studies. bioRxiv.

Deng, Q., Ramsköld, D., Reinius, B., and Sandberg, R. (2014). Single-cell RNA-seq reveals dynamic, random monoallelic gene expression in mammalian cells. Science, 343(6167), 193–196.

Devriendt, S., Antonio, K., Reynkens, T., and Verbelen, R. (2021). Sparse regression with multi-type regularized feature modeling. Insurance: Mathematics and Economics, 96, 248–261.

Edsgärd, D., Iglesias, M. J., Reilly, S.-J., Hamsten, A., Tornvall, P., Odeberg, J., and Emanuelsson, O. (2016). Geneiase: Detection of condition-dependent and static allele-specific expression from rna-seq data without haplotype information. Scientific Reports, 6(1), 21134.

Efron, B. and Morris, C. (1975). Data analysis using stein’s estimator and its generalizations. Journal of the American Statistical Association, 70(350), 311–319.

Fan, J., Wang, X., Xiao, R., and Li, M. (2021). Detecting cell-type-specific allelic expression imbalance by integrative analysis of bulk and single-cell rna sequencing data. PLoS Genetics, 17(3), e1009080.

Findley, A. S., Monziani, A., Richards, A. L., Rhodes, K., Ward, M. C., Kalita, C. A., Alazizi, A., Pazokitoroudi, A., Sankararaman, S., Wen, X., and et al. (2021). Functional dynamic genetic effects on gene regulation are specific to particular cell types and environmental conditions. eLife, 10.

Gutierrez-Arcelus, M., Baglaenko, Y., Arora, J., Hannes, S., Luo, Y., Amariuta, T., Teslovich, N., Rao, D. A., Ermann, J., Jonsson, A. H., et al. (2020). Allele-specific expression changes dynamically during T cell activation in HLA and other autoimmune loci. Nature genetics, 52(3), 247–253.

Hagemann-Jensen, M., Ziegenhain, C., Chen, P., Ramskold, D., Hendriks, G.-J., Larsson, A. J., Faridani, O. R., and Sandberg, R. (2020). Single-cell RNA counting at allele and isoform resolution using Smart-seq3. Nature Biotechnology, 38(6), 708–714.

Hannan, E. (1982). Spectral Analysis and Time Series-Priestley, MB. Metrika, 29, 212–212.

Hastie, T. J. (1992). Generalized additive models. In J. M. Chambers and T. J. Hastie, editors, Statistical Models in S, chapter 7. Wadsworth and Brooks/Cole, Pacific Grove, California.

Heinen, T., Secchia, S., Reddington, J., Zhao, B., Furlong, E., and Stegle, O. (2021). scDALI: Modelling allelic heterogeneity of DNA accessibility in single-cells reveals context-specific genetic regulation. bioRxiv.

Heinz, S., Romanoski, C. E., Benner, C., and Glass, C. K. (2015). The selection and function of cell type-specific enhancers. Nature Reviews. Molecular Cell Biology, 16(3), 144–154.

Hirai, H., Karian, P., and Kikyo, N. (2011). Regulation of embryonic stem cell self-renewal and pluripotency by leukaemia inhibitory factor. Biochemical Journal, 438(1), 11–23.

Hofling, H., Binder, H., and Schumacher, M. (2010). A coordinate-wise optimization algorithm for the Fused Lasso. arXiv preprint arXiv:1011.6f09.

Huber, W., Carey, V. J., Gentleman, R., Anders, S., Carlson, M., Carvalho, B. S., Bravo, H. C., Davis, S., Gatto, L., Girke, T., Gottardo, R., Hahne, F., Hansen, K. D., Irizarry, R. A., Lawrence, M., Love, M. I., MacDonald, J., Obenchain, V., Olés, A. K., Pagès, H., Reyes, A., Shannon, P., Smyth, G. K., Tenenbaum, D., Waldron, L., and Morgan, M. (2015). Orchestrating high-throughput genomic analysis with bioconductor. Nature Methods, 12(2), 115–121.

Jiang, Y., Zhang, N. R., and Li, M. (2017). SCALE: modeling allele-specific gene expression by single-cell RNA sequencing. Genome biology, 18(1), 1–15.

Khansefid, M., Pryce, J. E., Bolormaa, S., Chen, Y., Millen, C. A., Chamberlain, A. J., Vander Jagt, C. J., and Goddard, M. E. (2018). Comparing allele specific expression and local expression quantitative trait loci and the influence of gene expression on complex trait variation in cattle. BMC genomics, 19(1), 1–18.

Larsson, A. J., Johnsson, P., Hagemann-Jensen, M., Hartmanis, L., Faridani, O. R., Reinius, B., Segerstolpe, Å., Rivera, C. M., Ren, B., and Sandberg, R. (2019). Genomic encoding of transcriptional burst kinetics. Nature, 565(7738), 251–254.

Lawrence, M., Huber, W., Pagès, H., Aboyoun, P., Carlson, M., Gentleman, R., Morgan, M. T., and Carey, V. J. (2013). Software for computing and annotating genomic ranges. PLOS Computational Biology, 9(8), 1–10.

Picelli, S., Faridani, O. R., Björklund, Å. K., Winberg, G., Sagasser, S., and Sandberg, R. (2014). Full-length rna-seq from single cells using smart-seq2. Nature protocols, 9(1), 171–181.

Santoni, F. A., Stamoulis, G., Garieri, M., Falconnet, E., Ribaux, P., Borel, C., and Antonarakis, S. E. (2017). Detection of imprinted genes by single-cell allele-specific gene expression. The American Journal of Human Genetics, 100(3), 444–453.

Skelly, D. A., Johansson, M., Madeoy, J., Wakefield, J., and Akey, J. M. (2011). A powerful and flexible statistical framework for testing hypotheses of allele-specific gene expression from rna-seq data. Genome Research, 21(10), 1728–1737.

Stephens, M. (2017). False discovery rates: a new deal. Biostatistics, 18(2), 275–294.

Sun, W. (2012). A statistical framework for eQTL mapping using RNA-seq data. Biometrics, 68(1), 1–11.

Tian, L., Jabbari, J. S., Thijssen, R., Gouil, Q., Amarasinghe, S. L., Kariyawasam, H., Su, S., Dong, X., Law, C. W., Lucattini, A., et al. (2020). Comprehensive characterization of single cell full-length isoforms in human and mouse with long-read sequencing. bioRxiv.

Van Der Wijst, M. G., Brugge, H., de Vries, D. H., Deelen, P., Swertz, M. A., and Franke, L. (2018). Single-cell RNA sequencing identifies celltype-specific cis-eQTLs and co-expression QTLs. Nature genetics, 50(4), 493–497.

Vigorito, E., Lin, W.-Y., Starr, C., Kirk, P. D. W., White, S. R., and Wallace, C. (2021). Detection of quantitative trait loci from rna-seq data with or without genotypes using baseqtl. Nature Computational Science, 1(6), 421–432.

Wills, Q. F., Livak, K. J., Tipping, A. J., Enver, T., Goldson, A. J., Sexton, D. W., and Holmes, C. (2013). Single-cell gene expression analysis reveals genetic associations masked in whole-tissue experiments. Nature biotechnology, 31(8), 748–752.

Young, M. D., Wakefield, M. J., Smyth, G. K., and Oshlack, A. (2010). Gene ontology analysis for rna-seq: accounting for selection bias. Genome biology, 11(2), 1–12.

Zhu, A., Ibrahim, J. G., and Love, M. I. (2018). Heavy-tailed prior distributions for sequence count data: removing the noise and preserving large differences. Bioinformatics, 35(12), 2084–2092.

Zitovsky, J. and Love, M. (2020). Fast effect size shrinkage software for beta-binomial models of allelic imbalance [version 2; peer review: 1 approved, 2 approved with reservations]. F1000Research, 8(2024).

