## Supplementary for "*Airpart*: Interpretable statistical models for analyzing allelic imbalance in single-cell datasets"

### 1 Supplementary Methods

#### 1.1 Gene clustering

The Gaussian Mixture Model (GMM) method implemented in *mclust* (Scrucca *et al.*, 2016) and the hierarchical clustering method implemented in *dynamicTreeCut* (Langfelder *et al.*, 2007) (both methods included as alternatives in *airpart*) were used in order to detect gene clusters $C_u$ for $u \in \{1, ..., U\}$ according to the allelic ratio where $U$ is the number of gene clusters. For both methods, instead of directly providing the observed allelic ratios, first dimensionality reduction was performed on the observed allelic ratios via principal component analysis (PCA), for speed concerns and noise reduction. The data were clustered in a reduced dimensional space of twice the number of cell types, which is set in advance, either via prior knowledge or clustering cells using total counts. The transformed data in the reduced PC space were provided to the gene clustering methods. Briefly, the model-based clustering determines the most likely number and assignment of genes to clusters according to geometric properties of the data (distribution, volume, and shape). Within the GMM, an Expectation-Maximization algorithm is used for maximum likelihood estimation. The best model was selected according to Bayes Information Criteria, where *mclust* was run with `modelNames="EII"` both in the main model and initialization, which assumes the clusters all have a spherical distribution with equal volume. This model type was found to provide the most coherent clusters in terms of DAI in exploratory analysis. As an alternative, the hierarchical clustering method was used. The *dynamicTreeCut* algorithm can automatically choose the best tree by analyzing the shape of the branches that iteratively decomposing and combining clusters until find a stable solution. *dynamicTreeCut* was run with Manhattan distance and Ward agglomeration method. In the main Methods, for partition and allelic ratio inference, a single cluster is considered at a time. For notational simplicity, we omit index $u$ when referring to data from a gene cluster in the partitioning or allelic ratio inference sections.

#### 1.2 Data preprocessing

The following steps were applied to all four non-simulated datasets (Larsson *et al.* (2019), Deng *et al.* (2014), Combs and Fraser (2018), Gutierrez-Arcelus *et al.* (2020)), regardless of whether bulk or single cell. Cells/samples in which $> 5\%$ of the total counts (UMI counts for scRNA-seq) mapped to the mitochondrially encoded genes as well as those who exceed 3 median absolute deviations from the median detected genes were discarded. Genes that had 20 total reads in at least 25% cells/samples (Khansefid *et al.*, 2018) within each cell type or condition were kept for the Combs dataset, while five reads were used as a threshold for the Gutierrez-Arcelus dataset, and one read for the two single-cell datasets. As we were primarily interested in detecting *cis* genetic regulation (consistent imbalance relative to the genotypes),

*

and not imprinting or genome activation, the genes on the chromosome X were removed. Finally, only the gene set with largest ordered weighted mean allelic ratio difference $\geq 0.05$ that potentially exists differential allelic imbalance were modeled. For the Combs dataset, five reciprocal crosses (i.e. with either a *D. melanogaster* mother or a *D. simulans* mother) had different number of slices varying among 25-27, so we manually grouped the slices into 19 bins according to the correlation of consecutive slices across replicates. Three samples (a slice from a particular embryo) had relatively low correlation with other nearby slices and were dropped, and three samples were filtered out because of low total counts. Then, we performed a likelihood ratio test to detect genes with a significant parent-of-origin effect on allelic imbalance, and these were removed in order to focus on genetic effects. For data from Gutierrez-Arcelus, as signals from the same gene are tightly correlated and it would disrupt the inference if we dealt with each SNP independently, so we chose the SNP with largest total counts per gene. For polymorphic genes, such as genes in the HLA region, we note that we ignore the allelic complexity of these regions, by grouping alleles into two arbitrary groups (based on the SNP with the largest total count). This approach is not recommended for analysis of polymorphic genes. *airpart* only supports modeling of bi-allelic data (only two alleles present across all samples), and so this simplification was used for demonstration only. After exploratory data analysis of the allelic ratio, we determined there were individual differences in the temporal dynamics of allelic expression, and so we included a per-individual coefficient in the GFL and a per-individual offset in the allelic ratio estimation models. Finally, in order to focus on the most interesting patterns of allelic imbalance after T-cell stimulation, we used the variance of weighted allelic ratio mean to prioritize the top 43 most-varying genes for analysis for this dataset.

### 2 Supplementary Results

#### 2.1 Spatially varying allelic imbalance

We evaluated *airpart*'s performance on a spatial bulk dataset, where *Drosophila* F1 cross embryos had been sliced along the anterior-posterior axis and the RNA sequenced (Combs and Fraser, 2018). The analysis of this dataset had revealed a number of genes with spatially varying allelic imbalance, indicating that the two strains have diverged in the positioning of *cis* genetic regulation during development. We created a graph $\Gamma$ with edges only between consecutive groups of slices, reflecting the positional nature of the data. After removing the genes having clear parent-of-origin expression patterns that we interpreted as due to maternal deposition and quality control, we had 222 genes remaining. After exploratory data analysis (gene clustering followed by examination of allelic ratio heatmaps), we chose to set a large number of gene clusters (60) because there were a lot of divergent across genes and 4 gene clusters were especially interested shown in Figure S5 and others in Figure S6.

Raw observed allelic ratio with loess regression was used as a reference for comparison of fitted values from different models (Figure S5A). We saw the overall trend as well as the small up-turn at the anterior position in gene *tutl* and *CG13868* were both successfully captured by *airpart*. All genes in this cluster have allelic imbalance in anterior and posterior regions, e.g *CG30015* rejected the null hypothesis and there exists allelic imbalance among grouped slices 1-2,3-4 and 18-19 with credible interval (0.316, 0.385), (0.408, 0.471), (0.216,0.293), respectively.

*scDALI* was used with the radial basis function (RBF) kernel to allow for cells with similar $\mathbf{x}$ value to have similar fitted allelic ratio. We also performed *airpart* without the grouping step and it accepted basis matrix with 5 degrees of freedom to generate natural cubic splines with hierarchical modeling on dispersion parameter. The fitted allelic ratios (Figure S5C,D) were more similar to the loess fitted line in Figure S5A. Given that the spatially varying allelic imbalance for these embryos are likely a result of continuous gradients in concentration of regulatory proteins along the anterior-posterior axis (Combs and Fraser, 2018), the radial basis function (RBF) kernel and splines are therefore a good modeling choice for this dataset.

For some genes, *scDALI* was sensitive to fluctuations in the count ratios (e.g. *tutl* middle positions) compared to *airpart*. One of the genes *aos*, with large allelic ratio variation, failed during modeling with *scDALI*. More examples of *airpart*'s fusion of consecutive grouped slices are provided in Figure S6.

### 3 Supplementary Tables

| | |
|---|---|
| $u \in \{1,\ldots,U\}$ | gene cluster index |
| $g \in \{1,\ldots,G_u\}$ | gene index in gene cluster $u$ |
| $i \in \{1,\ldots,I\}$ | cell index |
| $k \in \{1,\ldots,K\}$ | cell state index |
| $Y_{gi}$ | allelic counts for gene g and cell i |
| $r_{gi}$ | allelic ratio for gene g and cell i |
| $m_{gi}$ | total read counts for gene g and cell i |
| $x_i$ | cell states of cell i |
| $c_i$ | nuisance covariates of cell i |
| $\beta_k$ | coefficient for cell state $k$ and $(\beta_0, \beta_1, \ldots, \beta_K) = \boldsymbol{\beta}$ |
| $\boldsymbol{\gamma}$ | coefficient vector for nuisance covariates |
| $N_{\boldsymbol{\Gamma}}$ | the total number of edges in graph $\boldsymbol{\Gamma}$ |
| $n_k$ | the number of cells in cell state k |
| $\mathcal{S}[K,K]$ | similarity score matrix with dimension $K \times K$ |
| $\mathcal{S}'[K,K]$ | network adjacency matrix with dimension $K \times K$ |
| $N$ | the number of elements within gene cluster. $N = G \times I$ |
| $q$ | tuning parameter for non-parametric test |
| $K_q$ | number of groups derived from constructing adjacency matrix according to q threshold among the partition of K cell types |
| $\hat{\sigma}_e^2$ | biased estimator for the true variance in non-parametric test |
| $j \in \{1,\ldots,J\}$ | cell group index |
| $l \in \{1,\ldots,N_j\}$ | cells who fall into cell group $j$ index |
| $\phi_g$ | true gene-wise overdispersion parameter |
| $\psi_g$ | gene-wise overdispersion estimates |
| $\psi_0$ | global gene dispersion hyper-parameter |
| $D$ | sampling variance of dispersion estimates $\psi_g$ |
| $A$ | global dispersion parameter variance |
| $B$ | weighted shrinkage between gene-wise and global variance estimators |
| $\boldsymbol{\beta}_g'$ | allelic ratio coefficient vector for gene $g$ in hierarchical Bayesian modelling |
| $\hat{\boldsymbol{\mu}}$ | estimates of the center of $\boldsymbol{\beta}_g'$ |

Table S1: Notation for the *airpart* model.

| Assessment | Metric | # of cells / cell type | # of genes / gene cluster | True allelic ratio | # of iterations |
|---|---|---|---|---|---|
| Partition* | ARI | 40 | {5, 10, 20} | {0.95, 0.9, 0.85, 0.85, 0.7, 0.7, 0.65, 0.6, 0.5, 0.5} | 200 |
| Estimation | RMSE | {40,100} | 1 | {0.3, 0.3, 0.3 + d, 0.3 + d, 0.3+2d, 0.3+2d, 0.3 + d, 0.3+d}** | 400 |
| DAI testing | TP/FP | {40,100} | 1 | {0.5,0.5,0.5,0.5,0.5,0.5} vs {0.5,0.5,0.6,0.6,0.7,0.7} | 400 |

* $cnt \in \{5, 10, 20\}$ and $\phi \in \{3, 20\}$. For other assessments, *cnt* is fixed at 10 and $\phi$ is 20.
** $d \in \{0.05, 0.1, 0.2, 0.3\}$

Table S2: Summary of different simulations settings used in evaluating the partition, allelic ratio estimation, and DAI testing of *airpart* variants and *scDALI*.

| method | scDALI | | airpart.nogroup | | airpart.bin | | airpart.gau | | airpart.np | |
|---|---|---|---|---|---|---|---|---|---|---|
| Truth \ Test | no DAI | DAI | no DAI | DAI | no DAI | DAI | no DAI | DAI | no DAI | DAI |
| no DAI | 97.75 | 2.25 | 97.00 | 3.00 | 93.75 | 6.25 | 89.50 | 10.50 | 88.00 | 12.00 |
| DAI | 2.00 | 98.00 | 1.75 | 98.25 | 2.50 | 97.5 | 2.25 | 97.75 | 0.50 | 99.50 |

Table S3: Statistical significance testing of DAI at $n = 40$.

| method | scDALI | | airpart.nogroup | | airpart.bin | | airpart.gau | | airpart.np | |
|---|---|---|---|---|---|---|---|---|---|---|
| Truth \ Test | no DAI | DAI | no DAI | DAI | no DAI | DAI | no DAI | DAI | no DAI | DAI |
| no DAI | 97.25 | 2.75 | 96.75 | 3.25 | 98.25 | 1.75 | 85.25 | 14.75 | 88.25 | 11.75 |
| DAI | 0.00 | 100.00 | 0.00 | 100.00 | 0.00 | 100.00 | 0.00 | 100.00 | 0.00 | 100.00 |

Table S4: Statistical significance testing of DAI at $n = 100$.

### 4 Supplementary Figures

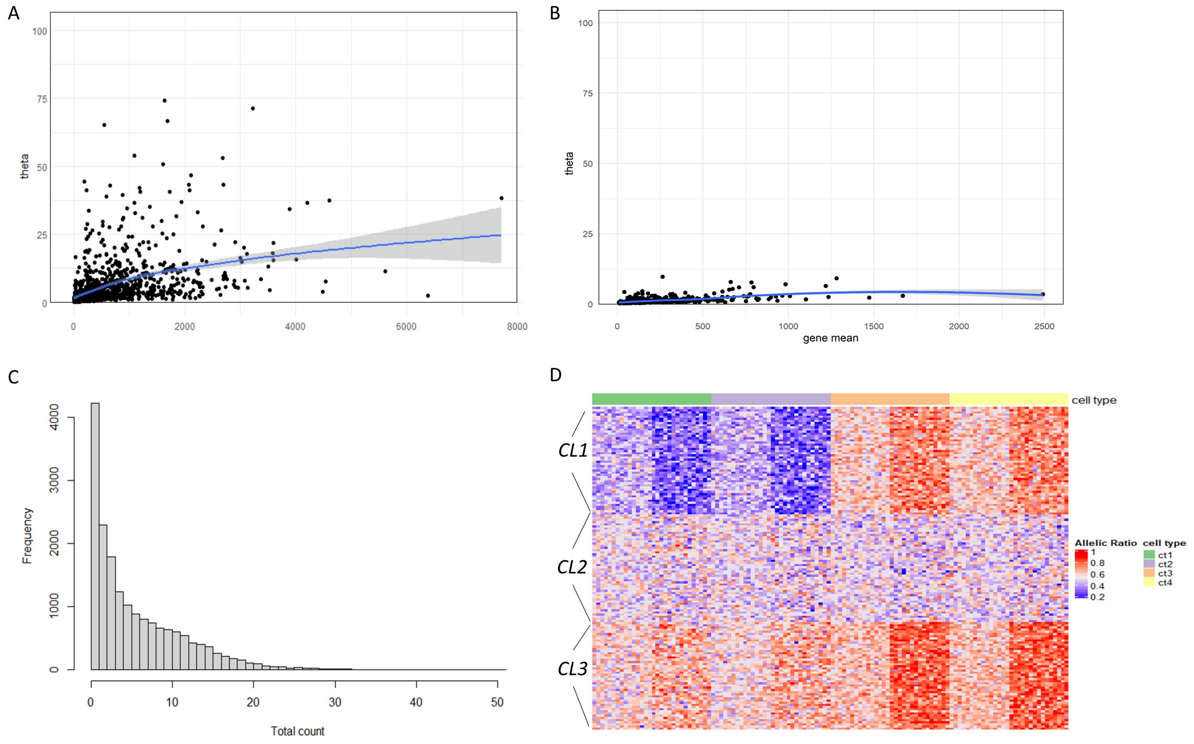


Figure S1: Statistical estimates for real and simulated single-cell datasets. A) Dispersion parameter estimates in Larsson's data. B) Dispersion parameter estimates in Deng's data. c) Simulated total count distribution with the higher mean count of NB = 10 and $\phi$=20. D) Gene clustering was performed to aggregate signal across genes with similar allelic ratio pattern. The true allelic ratios across 4 cell type of first 50 genes were {0.2,0.2,0.8,0.8}, the middle 50 genes were {0.5,0.5,0.5,0.5} and the last 50 genes were {0.7,0.7,0.9,0.9}.

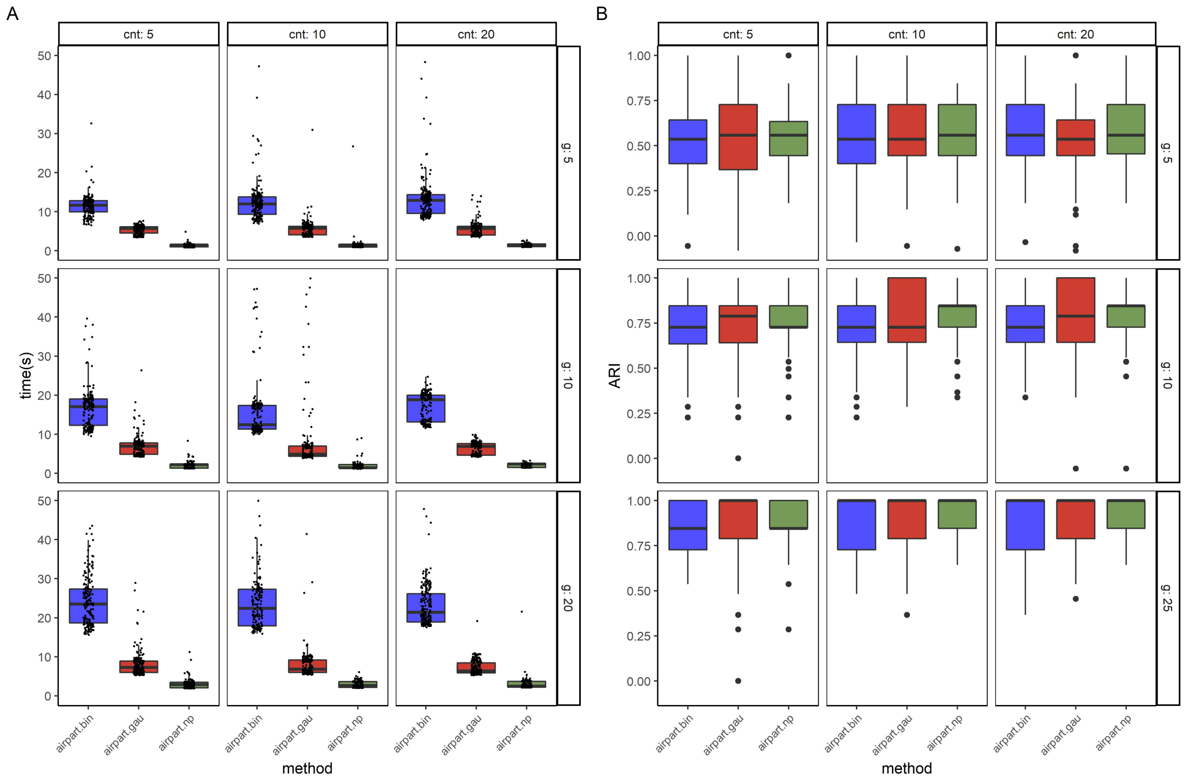


Figure S2: Performance comparison of airpart in running time and partition accuracy in highly dispersed simulation case A) Computation time of 1 cores among 200 iterations for the $\theta = 20$ and $n = 40$ simulation. B) Barplot of partition accuracy among 3 variants of *airpart* for $\theta = 3$ (over-dispersed allelic counts). The y axis is adjusted Rand index for 200 iterations. cnt: the higher mean count, g: number of genes within a gene cluster.

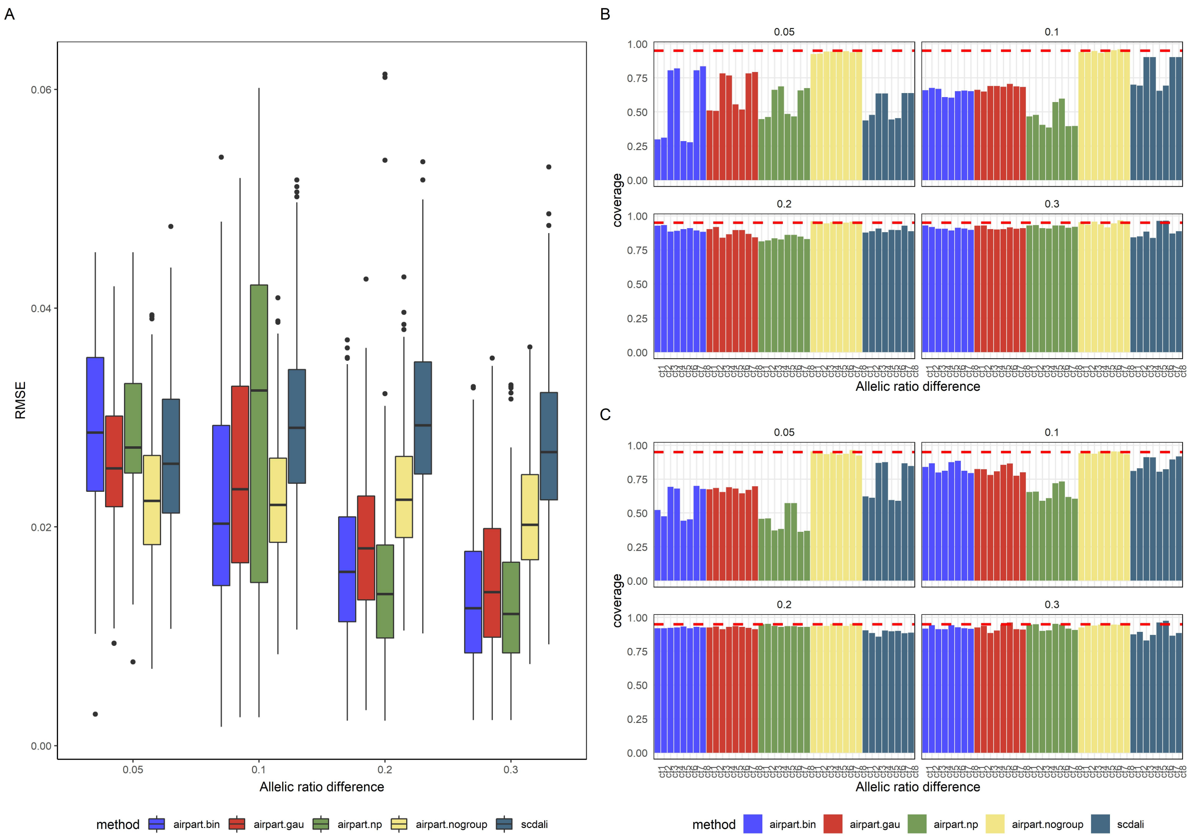


Figure S3: Performance comparison of *airpart* and *scDALI* on additional simulations. A) Boxplot shows RMSE of estimating allelic ratio on $n = 100$ among 400 iterations. Each gene has a U-shape pattern described in the Section 2.5 B) Barplot of 95% credible interval coverage over those observations in which the algorithms converged with n=40 setting on each cell type and each methods. C) Barplot of 95% credible interval coverage over those observations in which the algorithms converged with $n = 100$ setting on each cell type and each methods.

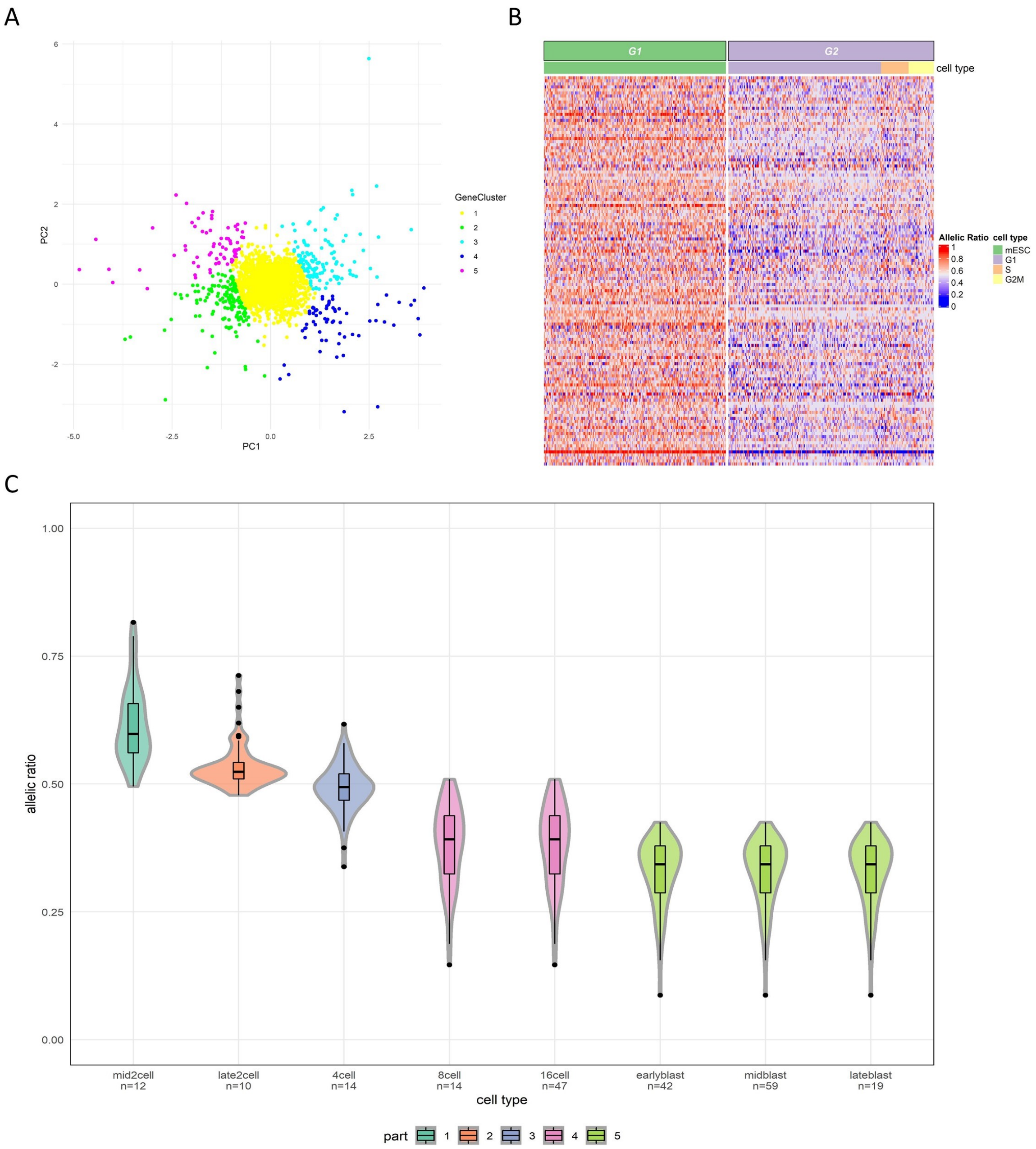


Figure S4: *airpart* result for Larsson's scRNA-seq data. (A) PCA plot of gene clustering step, (B) Heatmap of one gene cluster's allelic ratio with *airpart* cell type partition displayed on top, (C) Violin plot of allelic ratio estimates corresponds to Figure 3C using the Deng's dataset. Color represents different partition groups.

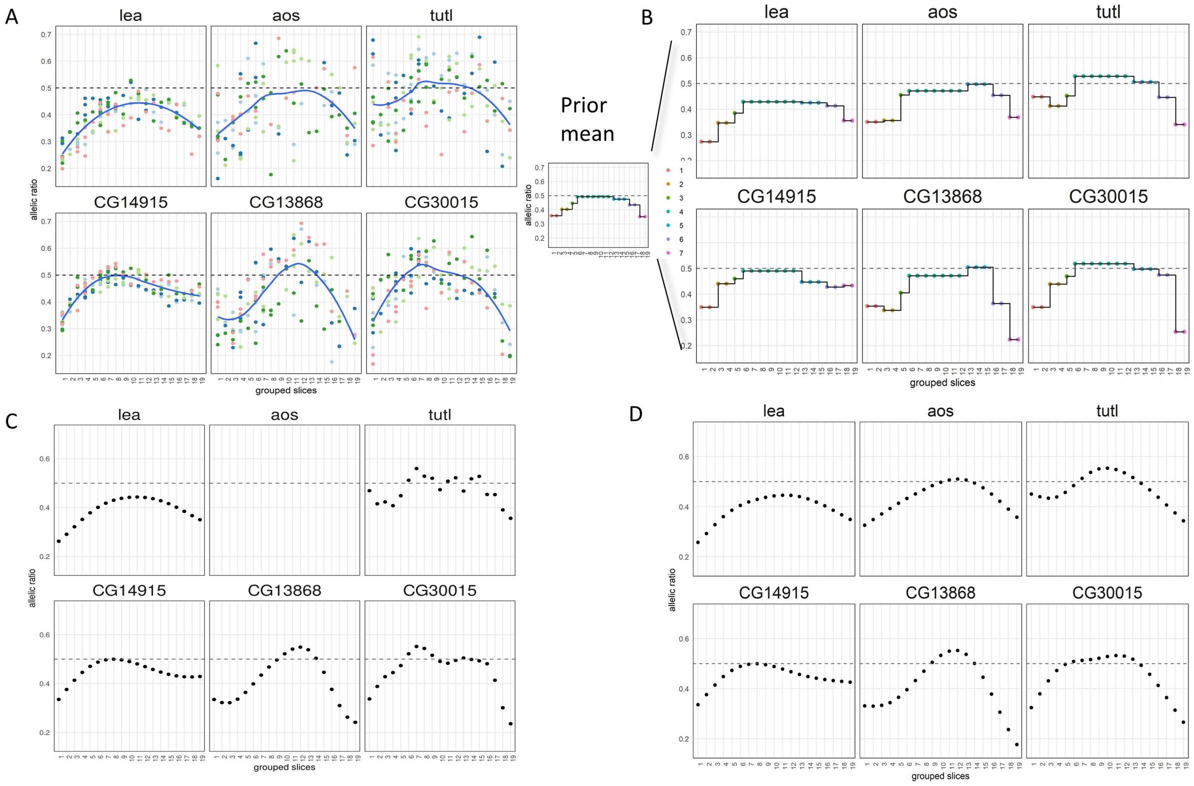


Figure S5: Evaluation of genes with differential allelic imbalance at different locations for Combs' data. x axis is group slices. For each gene, anterior is left and posterior is right with higher allelic ratio indicating D. simulans biased expression, and lower indicating D. melanogaster biased expression. A) Raw ratio with loess as smoothing blue line. Color represents different replicates. B) Estimates using *airpart*. C) Estimates using *scDALI*. D)Estimates using *airpart.no group*

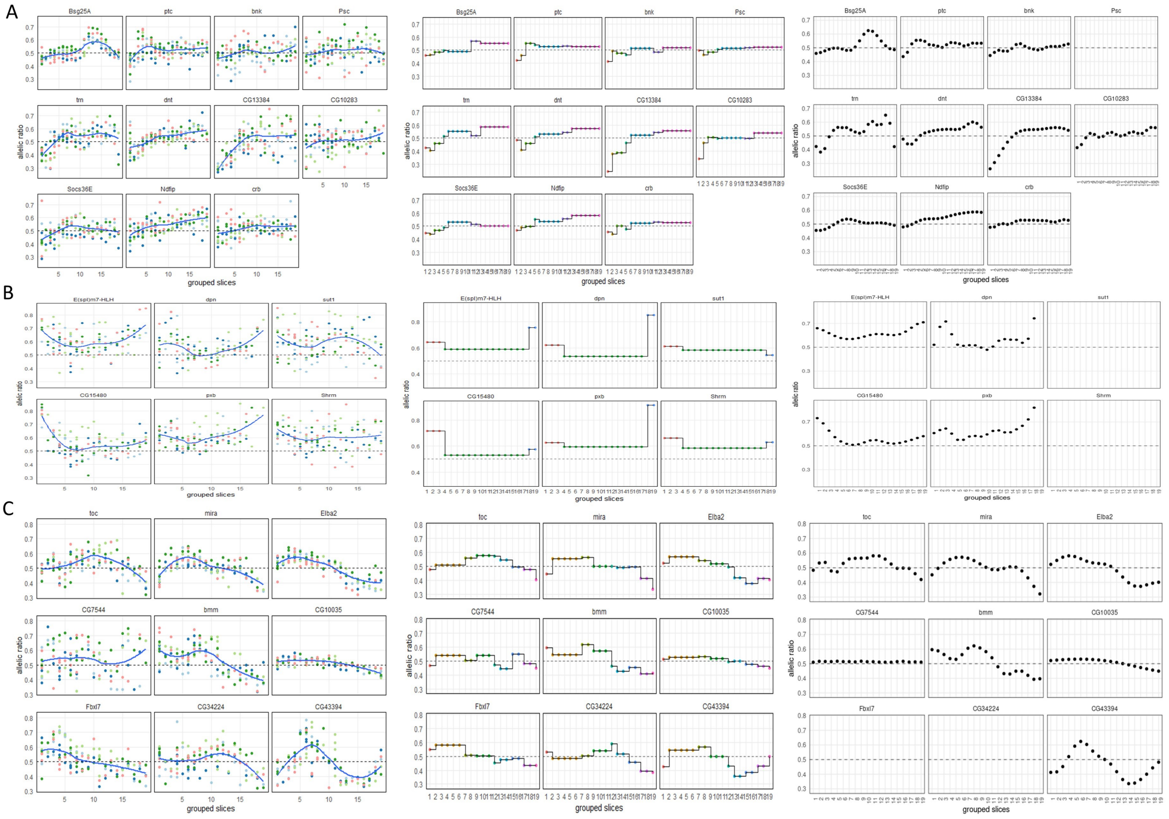

Figure S6: Three additional gene clusters showing *airpart* results for Combs' data. x axis is group slices. For each gene, anterior is left and posterior is right with higher allelic ratio indicating D. simulans biased expression, and lower indicating D. melanogaster biased expression.

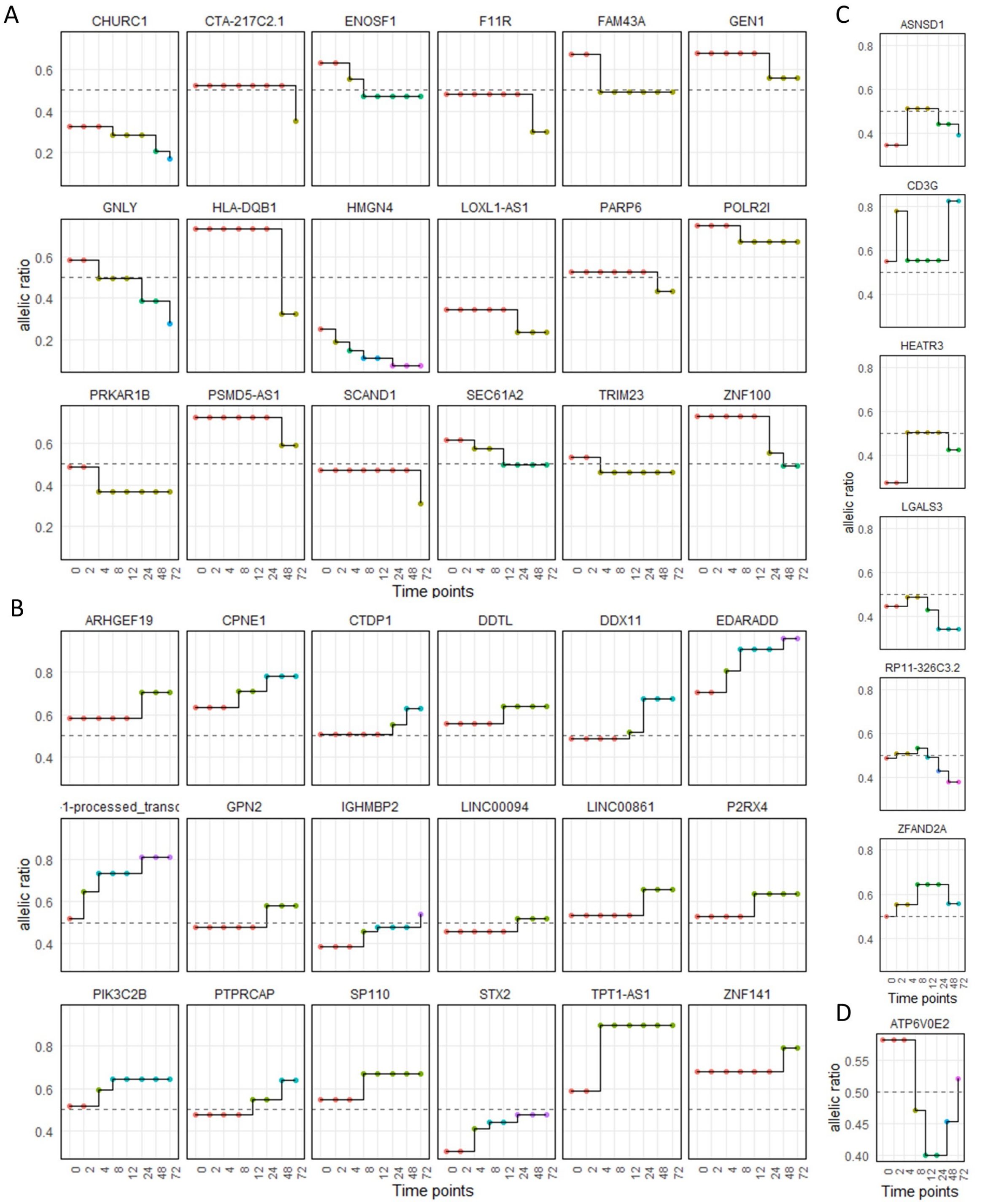


Figure S7: Complete step plot of ASE for top 43 genes with dynamic ASE. Genes are listed in the alphabetic order. A) decreasing ASE trend. B) increasing trend. C) up-down trend and D) down-up trend.

### References


Combs, P. A. and Fraser, H. B. (2018). Spatially varying cis-regulatory divergence in Drosophila embryos elucidates cis-regulatory logic. *PLOS Genetics*, **14**(11), 1–23.

Deng, Q., Ramsköld, D., Reinius, B., and Sandberg, R. (2014). Single-cell RNA-seq reveals dynamic, random monoallelic gene expression in mammalian cells. *Science*, **343**(6167), 193–196.

Gutierrez-Arcelus, M., Baglaenko, Y., Arora, J., Hannes, S., Luo, Y., Amariuta, T., Teslovich, N., Rao, D. A., Ermann, J., Jonsson, A. H., *et al.* (2020). Allele-specific expression changes dynamically during T cell activation in HLA and other autoimmune loci. *Nature genetics*, **52**(3), 247–253.

Khansefid, M., Pryce, J. E., Bolormaa, S., Chen, Y., Millen, C. A., Chamberlain, A. J., Vander Jagt, C. J., and Goddard, M. E. (2018). Comparing allele specific expression and local expression quantitative trait loci and the influence of gene expression on complex trait variation in cattle. *BMC genomics*, **19**(1), 1–18.

Langfelder, P., Zhang, B., and Horvath, S. (2007). Defining clusters from a hierarchical cluster tree: the Dynamic Tree Cut package for R. *Bioinformatics*, **24**(5), 719–720.

Larsson, A. J., Johnsson, P., Hagemann-Jensen, M., Hartmanis, L., Faridani, O. R., Reinius, B., Segerstolpe, Å., Rivera, C. M., Ren, B., and Sandberg, R. (2019). Genomic encoding of transcriptional burst kinetics. *Nature*, **565**(7738), 251–254.

Scrucca, L., Fop, M., Murphy, T. B., and Raftery, A. E. (2016). mclust 5: clustering, classification and density estimation using Gaussian finite mixture models. *The R journal*, **8**(1), 289.
